# The urea cycle is transcriptionally controlled by hypoxia-inducible factors

**DOI:** 10.1101/2021.01.25.428152

**Authors:** Charandeep Singh, Andrew Benos, Allison Grenell, Vincent Tran, Demiana Hanna, Bela Anand-Apte, Henri Brunengraber, Jonathan E. Sears

## Abstract

Here, we demonstrate transcriptional regulation of urea cycle genes CPS1 and ARG1 by hypoxia-inducible factors (HIFs) and demonstrate a hepatic HIF dependent increase in urea cycle activity.

## Introduction

The urea cycle is a liver specific pathway responsible for the removal of ammonia in ureotelic and uricotelic animals. Recently, recycling of ammonia has been shown to accelerate the proliferation rate of cancer cells (1, 2). Molecular nitrogen is not directly used by enzymes in the human body; we rather acquire our nitrogen from ammonia fixed into amino acids and proteins by microbes and legumes. Therefore, to preserve nitrogen for biosynthetic needs, the metabolism of nitrogenous compounds is very tightly regulated.

Conversion of ammonia to urea or orotate in mammals takes place in the liver, where it is first converted to carbamoyl phosphate. Enzymatic conversion of ammonia into carbamoyl phosphate, which is synthesized by Carbamoyl Phosphate Synthetase 1 (CPS1), is a rate-limiting step of the urea cycle. Carbamoyl phosphate has two fates, either to enter the urea cycle or to produce orotate-which is necessary for nucleotide synthesis and to transfer electrons to newly discovered electron transport chain component DHODH(3). If carbamoyl phosphate enters the urea cycle, arginine is synthesized from ammonia, which is converted either to urea or nitric oxide. The enzyme Arginase 1 (ARG1), a rate limiting enzyme in ureagenesis, in the liver converts arginine into urea and ornithine. The liver specific urea cycle is depicted in Figure 1. In addition to urea cycle, Arginase 2 (ARG2) activity is a rate limiting step in the polyamine synthesis and controls proliferation in some cell types (4). The liver specific urea cycle is depicted in Figure 1.

**Figure 1.**
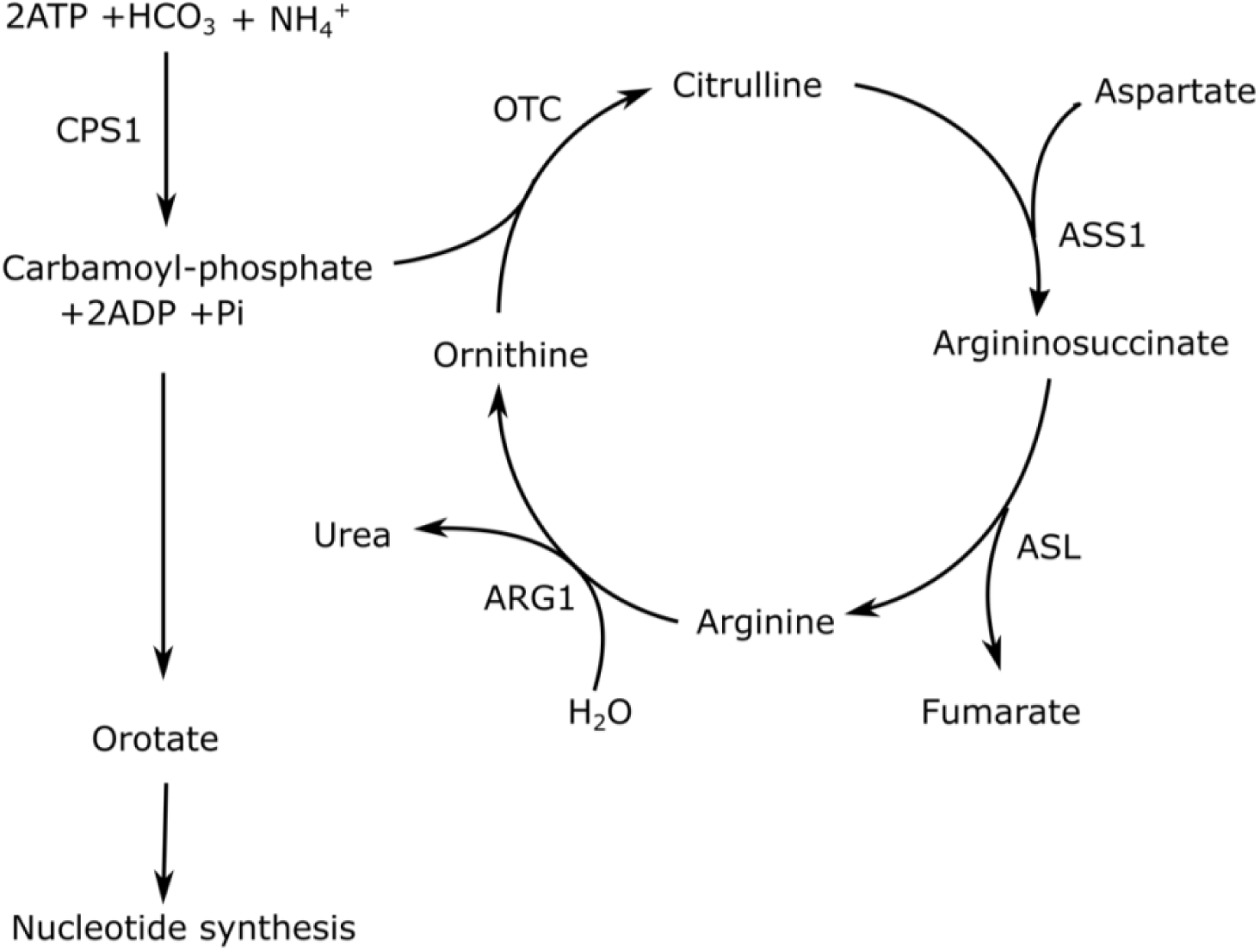
Urea cycle. Depicted are the liver specific urea cycle pathway enzymes and metabolites. Earlier studies have highlighted that the first enzyme of the pathway, CPS1, is only present in the liver or primary hepatocytes. CPS1 is the rate liming enzyme in the urea cycle pathway and it controls production of not just urea but also precursors for nucleotide synthesis and substrate for newly discovered electron transport chain component DHODH. Out of all the enzymes depicted in the picture, only CPS1 and ARG1 are rate limiting for urea cycle activity, with former playing roles more than merely in urea production and later controlling flux distribution between nitric oxide vs. ureagenesis.

Glucagon, glucocorticoids and insulin are known hormonal regulators of most urea cycle enzymes (5). Glucagon and glucocorticoids have transcriptional control of ASS1, ASL, and ARG1, stabilization of CPS1 mRNA and ARG1, and the protein stabilization of ornithine transcarbamylase (OTC) (6, 7). Additionally, transcriptional activators that control CPS1 expression are C/EBP and Hepatocyte Nuclear Factor 3 (HNF3-β) (5, 8, 9). The only known repressor of urea cycle genes CPS1, OTC and ARG1 transcriptionally is p53 (10).

## Results and Discussion

We previously demonstrated that a HIF prolyl hydroxylase inhibitor (HIF PHi)/stabilizer of HIF upregulated urea cycle metabolites in hyperoxic mice (11). To establish a direct connection of HIFs to urea cycle enzymes, we did qPCR analyses of a few urea cycle genes in primary hepatocytes isolated from 12 week old. We found increased expression of CPS1 when cells were treated with FG-4592, a HIF PHi (Fig. 2a). We also found increased expression of ARG1 when cells were treated with FG-4592 (Fig. 2b).

**Figure 2.**
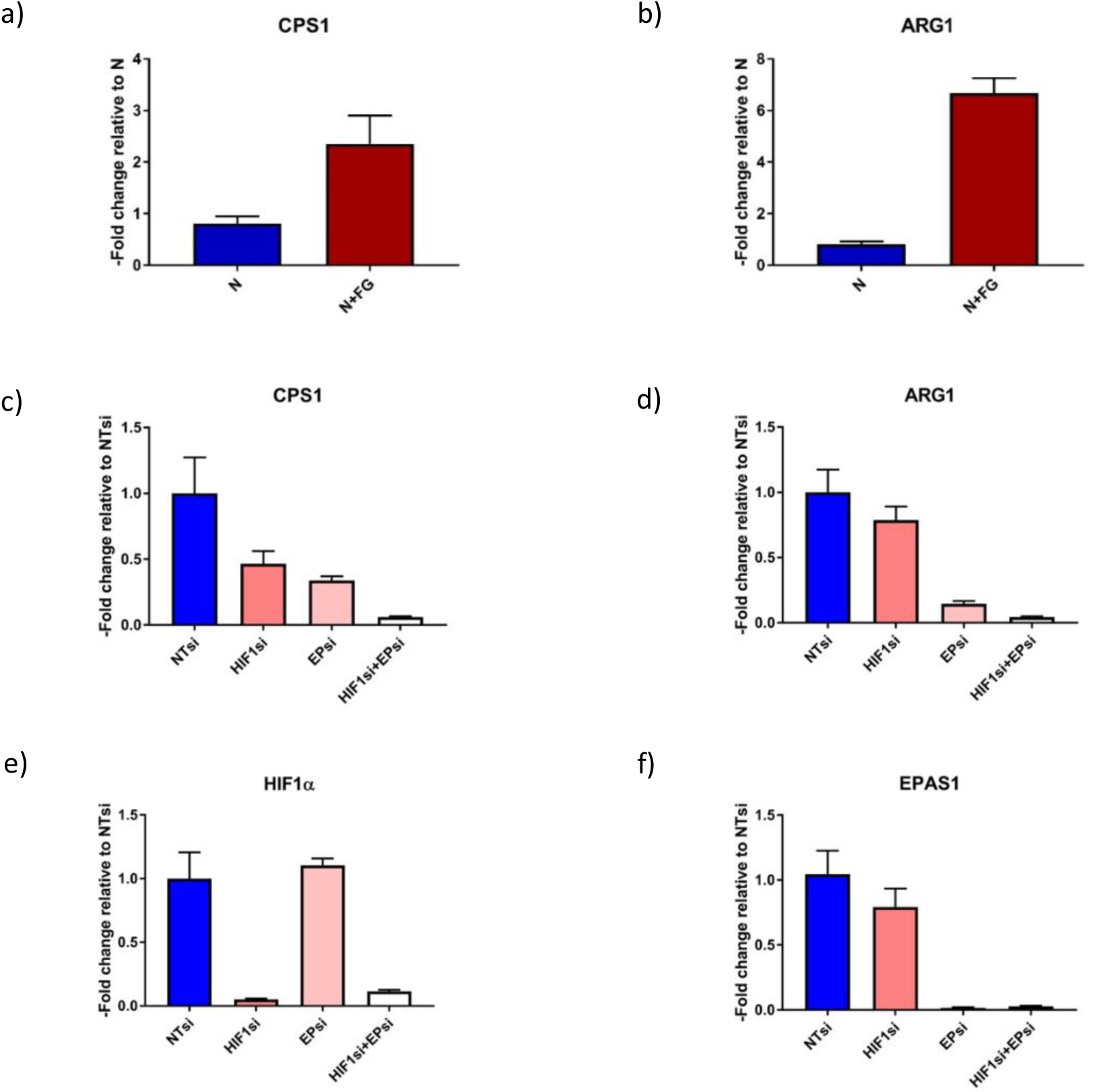
HIF controls expression of CPS1 and ARG1. Expression levels of a) CPS1 and b) ARG1 were found to be increased in response to FG-4592 treatment. Expression levels of c) CPS1 and d) ARG1 were downregulated in hepatocytes treated with siRNA against HIF1α or EPAS1. We measured expression of e) HIF1α and f) EPAS1 in hepatocytes treated with siRNAs against HIF1α and EPAS1 to confirm these siRNAs were specific for each HIF isomer. Primary hepatocytes were cultured in normoxia or hyperoxia for 24h with or without FG-4592 treatment. Gene expression fold change was calculated using 2^-ΔΔCt^ method. Gene expression was normalized to β-actin (ACTB) which served as the housekeeping control. Conditions: N, Normoxia; N + FG, Normoxia + FG-4592 treated. Legends: NTsi, non-targeting siRNA; HIF1si, HIF1α siRNA; EPsi, EPAS1 siRNA; HIF1si+EPsi, HIF1α siRNA+ EPAS1 siRNA.

We next tested to see if siRNA based knockdown (KD) of HIF1α and EPAS1 could modulate the expression levels of CPS1 and ARG1. Expression levels of CPS1 and ARG1 were decreased in the HIF1α and EPAS1 KD conditions, however, there were still some remaining transcripts in single KDs. We found a synergistic effect of HIF1 and EPAS1 KD on transcriptional activity of these genes. In the double KD, expression of CPS1 and ARG1 was near negligible (Figs. 2c and d). We measured the expression levels of HIF1α to ascertain that the synergistic effect was not due to cross-reactivity of HIF1α and EPAS1 siRNAs. We rather saw normal levels of HIF1α in the EPAS1 KD cells and near normal levels of EPAS1 in HIF1α KD (Figs. 2e and f).

To further confirm our findings at metabolic levels *in vivo*, we injected [^15^N_1_] ammonium chloride intraperitoneally in P12 mice that were treated with FG-4592 6h before ammonium chloride injections to stabilize HIF before measuring incorporation of labeled nitrogen. After injecting the [^15^N_1_] labeled ammonium chloride, livers from mice were dissected at 5, 8 and 15 minutes. FG-4592 injected animals had higher incorporation of labeled nitrogen from [^15^N_1_] ammonium into M1 labeled urea in the treated group versus in the controls (Figs. 3a, b and d). Liver Citrulline labeling was measured following [^15^N_1_] ammonium chloride injections and found to be higher in the FG-4592 treated animals (Fig. 3c). Plasma Arginine and Citrulline labeling were also investigated, and the maximal activity, measured as maximal initial enzyme velocity in the presence of excess substrate, in FG-4592 treated animals was higher than in control animals (Fig. 3e and f) indicating higher flux through the urea cycle.

**Figure 3.**
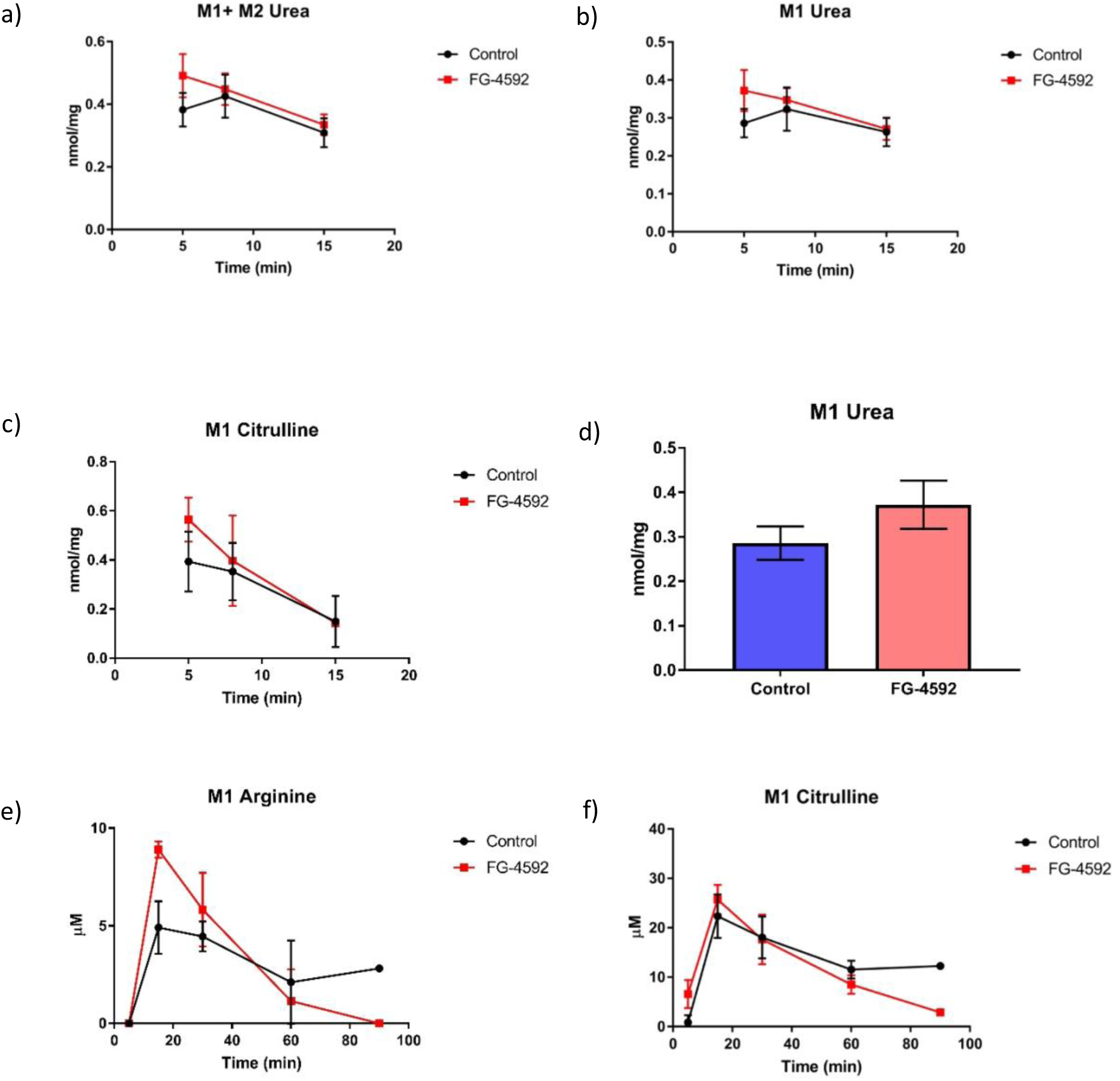
HIF stabilization by FG-4592 increases urea cycle flux *in vivo*. The absolute rates of liver specific (a) M1+M2 urea, (b) M1 urea, (c) M1 citrulline, and (d) histogram representation of M1 Urea at 5 min. Error bars represent standard deviations. (n=3, 5 min control; n=4, 8 min control; n=3, 15 min control; n=4, 5 min FG-4592; n=4, 8 min FG-4592; n=3, 15 min FG-4592). M1 Arginine time course for mouse plasma following injection of [^15^N_1_] ammonium chloride tracer (n=3 for 5 min, 15 min, and 30 min; n=2 for 60 min and 90 min) with and without systemic HIF stabilization. f) M1 Citrulline time course for mouse plasma following injection of [^15^N_1_] ammonium chloride tracer with and without (Figs 3e-f, n=3 for 5 min, 15 min, and 30 min; n=2 for 60 min and 90 min).

Known and unknown regulators of the urea cycle are depicted in Figure 4. Regulation by hormones occurs through the modulation of transcriptional regulation, mRNA stability, and protein stability (6). On the protein levels, post translational modification of Sirt3 and Sirt5 have been linked to deacetylation of urea cycle enzymes—thereby upregulating urea cycle activity (12, 13).. Li et. al. reported that p53 repressed expression of urea cycle genes CPS1, OTC and ARG1 (10) and also found increased expression of urea cycle genes in colon cancer cells, which implies a connection between hypoxia and urea cycle activity (10). Others and we have linked the urea cycle to the mouse model of oxygen induced retinopathy, which has alternating environments of hyperoxia and hypoxia (11, 14). Here, we definitively demonstrate that both mitochondrial CPS1 and cytosolic Arg1 are upregulated by HIF, providing global regulation of the urea cycle. We speculate that there are two ways HIFs can activate urea cycle gene expression: either by directly binding to hypoxia response elements on these genes or by repression of p53 activity which indirectly will activate expression of urea cycle genes.

**Figure 4.**
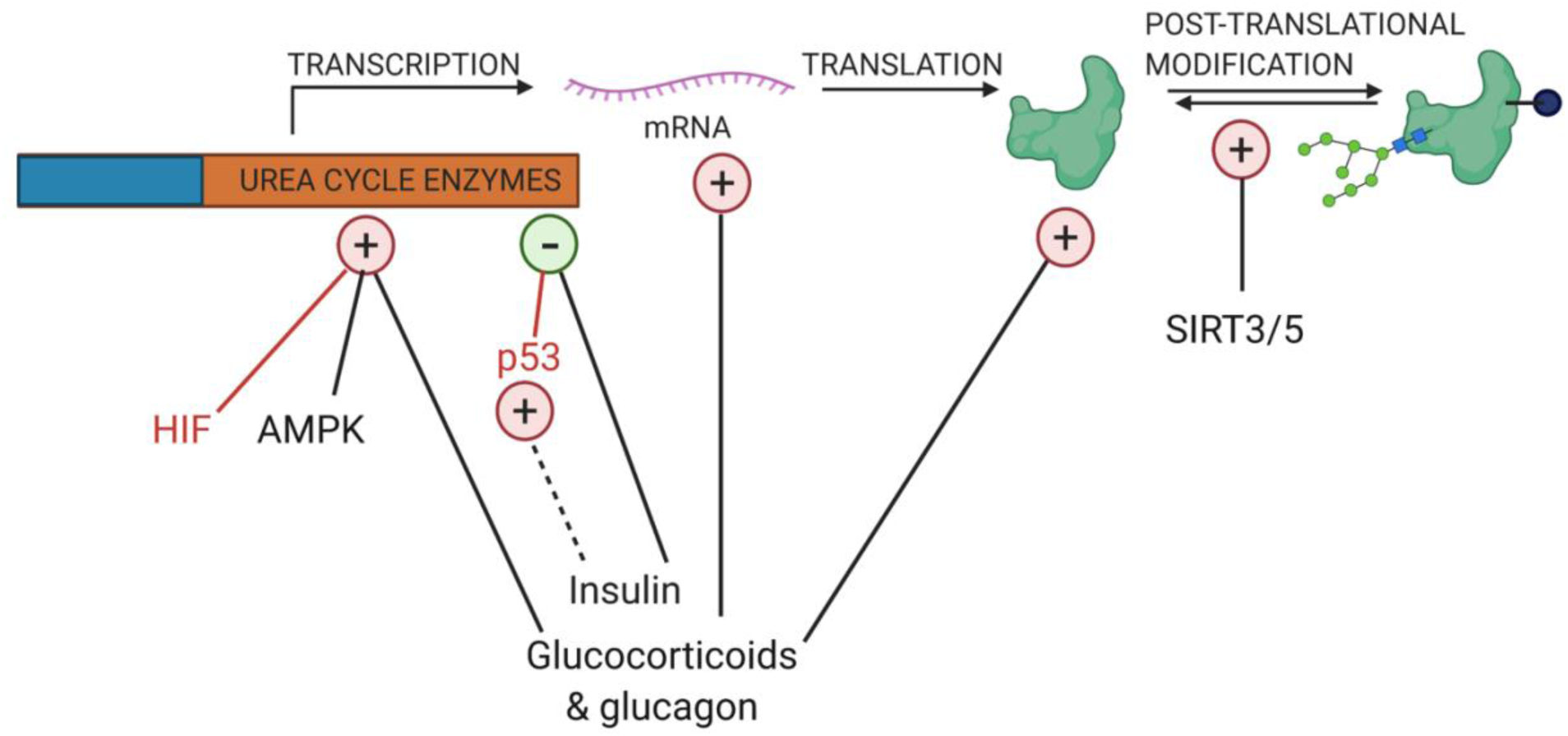
Known and unknown regulators of the urea cycle. Glucagon and glucocorticoids upregulate activity of urea cycle, whereas, insulin downregulates urea cycle activity. Sirt3/5 upregulates urea cycle activity by posttranslational modification of only mitochondrial urea cycle enzymes.

## Methods

### [^15^N_1_] ammonium chloride labeling of urea cycle compounds *in vivo* (liver)

Animals were kept in normal room air until p12. On p-12, pups were given 10 mg/Kg of FG-4592 subcutaneously and then around 6 h post injection 50 µl of 2.5 mg/ml of [^15^N_1_] ammonium chloride intraperitoneally (0.5 µmol/gm of body weight). Following the injection of ammonium chloride tracer, mice were anesthetized with isoflurane and then sacrificed at different time points. Livers were isolated and frozen on dry ice immediately.

### [^15^N_1_] ammonium chloride labeling of urea cycle compounds *in vivo* (plasma)

Animals were kept in normal room air until p10. On p-10, pups were given 10 mg/Kg of FG-4592 intraperitoneally and then around 4 h post injection 50 µl of 10 mg/ml of ammonium chloride subcutaneously. Post injection of ammonium chloride tracer, mice were sacrificed at different time points, 5min, 15min, 30min, 60min, and 90min. Blood was isolated from heart and added to heparin containing tubes and then centrifuged at 1020 x g for 20 min at 4°C.

Note: The doses of ammonium chloride were chosen close what has earlier been used in human subjects(15, 16). We recommend using lower dose if infusion experiment is deemed necessary, for example as used in Yang et. al.(17).

### Labeling data analysis

Areas of all mass isotopomers were extracted using masshunter data analysis software (Agilent). Data were corrected for naturally occurring istopes using Isocor software.

Simultaneously, total quantities were calculated first by obtaining ratios of total urea/internal standard and then quantified using external calibrations curve of urea/internal standard at different ranges. Following formula was used to calculate the quantities of each isotopmer

Total quantity (μ mol/mg of tissue) x isocor corrected fraction of M1 or M2 out of total value of 1 = Quantities of M1 or M2 labeled compounds in μ mol/mg of tissue

### Metabolite extraction from the livers

An extraction buffer consisting of 40 ml of 80% methanol and 400 μL of [^13^C_5_] ribitol (0.05 mg/ml) was prepared and kept at -20°C for use. Liver samples were removed from-80°C and thawed on ice. These samples were weighed, transferred into fresh tubes, and kept on ice. 250 μl of cold extraction buffer was added to each tube. The liver samples were homogenized using fresh disposable pestles then sonicated. The homogenized samples were then stored in -80°C freezer for 20 min to quench the metabolism. The samples were then removed and transferred into fresh tubes containing additional cold extraction buffer so that the final concentration of tissue in each sample was 25 μl (buffer)/mg (tissue). The samples were then vortexed and centrifuged at 3,000 x g for 10 min at ≤ 4°C. 500μL of supernatant was transferred into fresh tubes and dried over night at -4°C in a refrigerated centrivap concentrator (Labconco). The dried samples were then derivatized for GCMS measurement. All the samples were measured on EI-GCMS.

### Metabolite extraction from the plasma samples

Five microliters of plasma sample was added to 160 µl of -20°C cold acetonitrile and additional 35 µl of water was added to the samples. Samples were vortexed and immediately centrifuged at 15000 x g for 5 min at 4°C. Supernatant 150 µl was dried with 10ul of 0.05 mg/ml of [^13^C_5_] ribitol, overnight at -4°C in a refrigerated centrivap concentrator (Labconco).

### Derivatization of samples for GCMS measurement

Dried samples were removed from the centrivap concentrator (Labconco). 25 μL of Methoxyamine hydrochloride/pyridine (sigma Aldrich) solution (40 mg/ml) were added to each tube. The samples were mixed at 45°C at 1,000 rpm for 30 min on the ThermoMixer (Eppendorf). 75 μL of MSTFA + 1% TMCS (Thermo Scientific) were added to each tube and the tubes were mixed at 45°C at 1,000 rpm for a second 30-minute period. The samples were removed from the ThermoMixer and were allowed to cool to room temperature before centrifugation at 15,000 x g for 3 min (room temperature). Approximately 60 μl of supernatant were packed into GCMS vials, out of which 1ul was injected. GCMS method use for measurement was same as described earlier in Singh et. al. (2020) with slight changes: column used here was DB-5MSDG 40m x 250 µm × 0.25 µm (Agilent) and post-column re-equilibration step was removed (18).

### Cell culture, RNA extraction and cDNA preparation

Cells were seeded in 6-well plates in Williams E media at a staring density of 0.3 × 10_^6^_ cells per well. Plates were incubated in 5% CO_2_ incubator set at 37°C for one day. On the following day, some of the pates were treated with siRNA (details of siRNA treatment described below). On the third day, some of the plates were treated with FG-4592 (20ul of 1mg/ml FG-4592 added each well of the 6-well plate). Half of the plates were incubated in normoxia (21% O_2_, 5% CO_2_ at 37°C) and rest half in hyperoxia (75% O_2_, 5% CO_2_ at 37°C) incubator for 24h. Following the incubation, RNA was extracted from the cells using TriReagent. Media was aspirated from the plates, following which cells were washed with 1ml of normal saline and 1ml of TriReagent was added to each plate. Plates were incubated at room temperature for 5min, following which cell lysate was transferred to 1.5 ml tubes. Added 200 µl of chloroform to each tube and vortexed for 1min. The samples were then incubated samples at room temperature for 10 min and subsequently centrifuged at 12000 x g for 15 min at 4°C. The upper layer was transferred to fresh tubes containing 0.5 ml of isopropanol, mixed and then incubated at room temperature for 10 min. The samples were again centrifuged at 12000 x g for 10 min at 4°C and discarded the supernatants were discarded. The pellets were washed with 1ml of 75% ethanol followed by centrifugation at 7500 x g for 5min at 4°C.

Discarded supernatant and air dried pellets. RNA was dissolved in 20 µl of molecular biology grade water. RNA was converted to cDNA using Verso cDNA synthesis kit (Thermo Scientific) using anchored oligo-dT primers and following protocol provided with the kit.

### RT-qPCR

Two microliters of cDNA was added to each well of 96 well plate, followed by addition of 1x Radiant_^™^_ SYBR Green Lo-ROX (Alkali Scientific). Following qPCR primers were used

1. ARG1 Quantitech catalogue number QT00134288, 1x per reaction
2. HIF1α Quantitech catalogue number QT01039542, 1x per reaction Self-designed primers
3. CPS1, Forward 5’-GTGAAGGTCTTGGGCACATC-3’ Reverse 5’-TTCCACTGCAAAACTGGGTG-3’
4. β-actin Forward 5’-TACTCTGTGTGGATCGGTGG-3’ Reverse 5’-TCGTACTCCTGCTTGCTGAT-3’

Quantitech primers were used at 1x concentration per well. Self-designed primers were used at a final concentration of 1 µl of 10 µM forward and 1 µl of 10 µM reverse, per well. Volume of each well was made to 20 µl with molecular biology grade water. The qPCR program used was 50°C for 2 min, 95 °C for 10 min, 40 cycles at 95 °C for 00:15 min, 60 °C for 00:20 min, and 72 °C for 00:20 min with acquisition of fluorescence followed by a melting curve 95 °C for 00:15 min, 60 °C for 1:00min, 95 °C for 00:15 min.

### siRNA mediated knock-down

Following ON-TARGETplus non-targeting pool from Dharmacon (Horizon Discovery):

1. ON-TARGETplus Mouse Hif1a (15251) siRNA – SMARTpool, catalogue number L-040638-00-0005
2. ON-TARGETplus Mouse Epas1 (13819) siRNA – SMARTpool, catalogue number L-040635-01-0005
3. ON-TARGETplus Non-targeting Pool, catalogue number D-001810-10-05

5nmol of powdered siRNAs were dissolved in 100 µl of water and mixed well with pipetting up and down couple of times. siRNA solutions were mildly vortexed before use. 25 µl of siRNA solution was mixed with 625 µl of low-serum Optimem media. In a separate tube, 75 µl of lipofectamine 2000 (Invitrogen) was mixed with 625 µl of Optimem. Added siRNA/Optimem mixture to tube containing lipofectamine 2000/Optimem and incubated at room temperature for few mins. Media in the 6-well plates was changed with fresh low-serum Opimem, followed by the addition of 100 µl siRNA/lipofectamine/Optimem mixture per well. Plates were incubated for 5h in 5% CO2 incubator set at 37°C. Following which media was changed to William’s E media and further procedures were performed as described above.

## Acknowledgements

Grant Support: National Eye Institute (R01 EY024972 to JES; R01 EY026181 to BAA, R01 EY027083 to BAA, P30EY025585 to BAA); 5T32EY007157-19 grant to AG; Research to Prevent Blindness RPB1801 to JES. This work was supported in part by RPB Challenge grant and unrestricted grant from the Cleveland Eye Bank Foundation. We would like to thank Northern Ohio Alcohol Center for providing us freshly isolated hepatocytes, supported by grant P50AA024333 (PI Dr. Laura Nagy, Cleveland Clinic, Ohio).

Author Contributions: CS conceived, designed performed, analyzed, wrote the paper. AB performed experiments and data analysis, and edited the paper. AG performed experiments and edited the paper. VT performed experiments and analyzed data. DH performed experiments. BA conceived, designed and wrote the paper. HB conceived, designed and wrote the paper. JES conceived the experiments and wrote the paper.

## Data availability

All the mass spectrometry raw data are available online on metabolights, study id: MTBLS2221.

## Study approval

All experimental procedures involving live animals were conducted in accordance with the guidelines of the NIH’s Guide for the Care and Use of Laboratory Animals (National Academies Press, 2011) and approved by the Cleveland Clinic institutional animal care and use committee (IACUC, protocol 2016-1677). All animal experiments conformed to the rules and regulations of the Association of Research in Vision and Ophthalmology.

## Abbreviations

HIF PHi: hypoxia inducible factor-prolyl hydroxylase inhibition;
OIR: oxygen induced retinopathy;
CPS1: Carbamoyl phosphate synthetase I;
ARG1: Arginase 1; ARG2, Arginase 2;
OTC: Ornithine transcarbamylase;
HNF3-β: Hepatocyte nuclear factors;

